# Refining the genomic location of SNP variation affecting Atlantic salmon maturation timing at a key large-effect locus

**DOI:** 10.1101/2021.04.26.441431

**Authors:** Marion Sinclair-Waters, Nikolai Piavchenko, Annukka Ruokolainen, Tutku Aykanat, Jaakko Erkinaro, Craig R. Primmer

**Affiliations:** Organismal and Evolutionary Biology Research Programme, Faculty of Biological and Environmental Sciences University of Helsinki, Helsinki, Finland; Institute of Biotechnology, Helsinki Institute of Life Sciences, University of Helsinki, Helsinki, Finland; Natural Resources Institute Finland (LUKE), Oulu, Finland

## Abstract

Efforts to understand the genetic underpinnings of phenotypic variation are becoming more and more frequent in molecular ecology. Such efforts often lead to the identification of candidate regions showing signals of association and/or selection. These regions may contain multiple genes and therefore validation of which genes are actually responsible for the signal is required. In Atlantic salmon (*Salmo salar*), a large-effect locus for maturation timing, an ecologically important trait, occurs in a genomic region including two genes, *vgll3* and *akap11*, but data for clearly determining which of the genes (or both) contribute to the association have been lacking. Here, we take advantage of natural recombination events detected between the two candidate genes in a salmon broodstock to reduce linkage disequilibrium at the locus, and thus enabling delineation of the influence of variation at these two genes on maturation timing. By rearing 5895 males to maturation age, of which 81% had recombinant *vgll3*/*akap11* allelic combinations, we found that *vgll3* SNP variation was strongly associated with maturation timing, whereas there was little or no association between *akap11* SNP variation and maturation timing. These findings provide strong evidence supporting *vgll3* as the primary candidate gene in the chromosome 25 locus for influencing maturation timing. This will help guide future research for understanding the genetic processes controlling maturation timing. This also exemplifies the utility of natural recombinants to more precisely map causal variation underlying ecologically important phenotypic diversity.

## INTRODUCTION

The identification of genetic variation underlying phenotypic variation is a common goal in biology. A first step towards this goal is commonly a ‘genome scan’, where variation across the entire genome, or significant proportion of it, is scanned for signatures of selection and/or genotype-phenotype associations. When phenotype measurements are unavailable, or if there is no prior knowledge of adaptive phenotypes, genome scans identify loci potentially under selection via outlier testing (Pritchard et al. 2018; Kardos et al. 2015; Sinclair-Waters et al. 2017). Whereas when phenotypic measurements are available, genome scans can be used to search for associations between genetic and phenotypic variation (Barson et al. 2015; Johnston et al. 2014; 2011). When successful, signals of association and/or selection often lead to the identification of genomic regions including multiple candidate genes. In cases where a signal is particularly strong, a logical follow-up aim is to better validate which genes are actually linked to the signal. Such validation of candidate genes in model systems can be done via knock-outs (e.g. The International Mouse Knockout Consortium 2007; Varshney et al. 2013), and CRISPR (Sander and Joung 2014). Recently, candidate gene validation using CRISPR has been achieved in some free-living taxa such as butterflies (Livraghi et al. 2018; Concha et al. 2019; Woronik et al. 2019), sticklebacks (*Gasterosteus aculeatus*) (Wucherpfennig, Miller, and Kingsley 2019) and some crops (Rodríguez-Leal et al. 2017; Sedeek, Mahas, and Mahfouz 2019), but this is not yet feasible in many free-living species and likely will not enable testing of candidate variation in the wild. However traditional mapping approaches, where natural recombination events can be exploited to delineate the effects of linked genes, can be used when such natural recombinants are identified and where controlled crossing, followed by phenotypic assessment is feasible. Here, we use Atlantic salmon (*Salmo salar*) as a model system for how this approach can be applied to delineate the effects of linked genes at a locus associated with a trait of ecological relevance.

Atlantic salmon are an anadromous species that can spend one to seven years in freshwater, before migrating to the ocean where they can spend another one to five years before reaching maturation and returning to their natal rivers to spawn. Furthermore some (mostly) male individuals, known as mature parr, reach maturation in the freshwater environment without having migrated to sea. This age at maturity can vary both within and among populations, and contributes markedly to the diversity of life-history strategies of this species (Mobley et al. 2021; Erkinaro et al. 2019). Late maturation is associated with larger size, and therefore increased fecundity in females and greater reproductive success in males. Maturing at a later age, however, also increases the risk of mortality prior to reproduction (Mobley et al. 2020; Fleming and Einum 2011). Many loci with a variety of effect sizes are associated with Atlantic salmon maturation timing (Sinclair-Waters et al. 2020). One locus on chromosome 25, of particular interest due to its large effect size, explains close to 40% of the variation in age at maturity in both wild populations and aquaculture strains from Northern Europe (Barson et al. 2015; Ayllon et al. 2015). The SNP with the strongest association at this locus was located 7.9kb downstream of the *vgll3* gene and 45.4kb upstream of the *akap11* gene (Barson et al. 2015). In another association study using individual-level sequencing data capturing more sequence variation, Sinclair-Waters et al. (2021) also found that the SNP with the strongest association was located in the region between these two genes, however, slightly further downstream of *vgll3* (10.3kb) and closer to *akap11*. Additionally, two missense mutations occur within *vgll3* and one missense mutation occurs within *akap11*. Although not the most strongly associated SNPs with age at maturity, all three missense mutations showed a significant association signal in wild populations (Barson et al. 2015; Ayllon et al. 2015).

In addition to the strong association signals observed on chromosome 25, both *vgll3* and *akap11* are plausible candidates for influencing maturation timing given their reported functions. The *vgll3* gene, vestigial-like family member 3, is a transcription cofactor that inhibits adipogenesis and is associated with mouse weight and total fat mass (Halperin et al. 2013). In many species, including salmon, sufficient fat storage is needed to provide energy for maturation (Good and Davidson 2016), thus suggesting *vgll3* is a good candidate gene for Atlantic salmon maturation. Additionally, *VGLL3* is associated with age at maturity in humans (Cousminer et al. 2013; Day et al. 2017; Perry et al. 2014). The *akap11* gene encodes A-kinase anchoring protein 11. Evidence showing that A-kinase-anchoring proteins are expressed in testes during spermatogenesis and are important for sperm motility in humans (Reinton et al. 2000; Luconi et al. 2004) and mice (Miki et al. 2002) suggests that *akap11* may be important for sperm function and thus also a good candidate gene for involvement in Atlantic salmon maturation. Further, the expression patterns of *vgll3* and *akap11* have been shown to be correlated in various Atlantic salmon juvenile life history stages (Kurko et al. 2020). Both genes are plausible candidates for maturation and therefore determining whether the locus’ association with maturation timing is linked to *vgll3, akap11* or both genes is an important step for understanding the genetic process underlying variation in maturation timing in Atlantic salmon.

Here, we capitalize on the occurrence of a recombination event between the *vgll3* and *akap11* genes in a large number of individuals from a captive Atlantic salmon broodstock to delineate the effects of these two adjacent and physically linked genes, on maturation timing. Progeny from 16 independent families were bred using controlled crosses where at least one parent carried the recombinant alleles, and 5895 males were reared to maturation age. This allowed testing of whether the association of this chromosomal region with maturation is driven by SNP variation linked to *vgll3* or *akap11*, or a combination of both. The results provide greater resolution of the association signal at a known large-effect locus and help to narrow down the possible genomic location of causal variation underlying maturation timing in Atlantic salmon.

## METHODS

### Animal material

We reared 16 families using parental Atlantic salmon (*Salmo salar*) from a Neva river strain maintained at a Natural Resources Institute Finland hatchery in Laukaa, Finland (62°24’N, 25°57’E) (See Debes et al. 2019 for more broodstock details). Parents were chosen from a total of 702 broodstock individuals that had earlier been genotyped for 177 SNPs on Ion Torrent or Illumina (Miseq or Next-Seq) sequencing platforms as outlined in Aykanat et al. (2016). These SNPs included two missense SNPs in *vgll3*, the top-associated SNP from Barson et al. (Barson et al. 2015) located 7.9kb downstream of *vgll3*, and one missense SNP in *akap11*. We selected parents based on their *vgll3* and *akap11* genotypes that would maximize the proportion of offspring with a recombination event between the *vgll3* and *akap11* genes. For example, individuals carrying a haplotype with an *L* allele for *vgll3* and an *E* allele for *akap11*, or vice versa. We avoided crossing closely related individuals (those with grandparents in common) by using SNP-based pedigree reconstruction as in Debes et al. (2019). Additionally, we selected only parents that had the same genotype at the two *vgll3* missense mutations and top-associated non-coding SNP identified in Barson et al. (2015). From this point onwards, four character genotypes will be used to describe an individual’s genotype at the focal loci, *vgll3* and *akap11*. The first two characters indicate the genotype at the *vgll3* locus and the last two characters indicate the genotype at the *akap11* locus. The locus is indicated in subscript text after the genotype. Details of the 16 crosses are outlined in Table 1 (Supplementary Table 1).

**Table 1.** Description of the six types of crosses used including parental genotypes, number of families per cross type and proportions of offspring genotypes. The first and last two alleles listed indicate the *vgll3* and *akap11* genotypes, respectively, where *E* and *L* are the alleles found to be associated with earlier and later maturation, respectively, in Barson et al. (2015).

| Cross type | # of families | Offspring genotypes |
| --- | --- | --- |
| *EL<sub>vgll3</sub>EE<sub>akap11</sub>* × *EL<sub>vgll3</sub>EL<sub>akap11</sub>* | 3 | ~25% *EE<sub>vgll3</sub>EE<sub>akap11</sub>*, ~25% *EL<sub>vgll3</sub>EE<sub>akap11</sub>*,<br>~25% *EL<sub>vgll3</sub>EL<sub>akap11</sub>*, ~25% *LL<sub>vgll3</sub>EL<sub>akap11</sub>* |
| *LL<sub>vgll3</sub>EL<sub>akap11</sub>* × *EL<sub>vgll3</sub>EL<sub>akap11</sub>* | 6 | ~25% *EL<sub>vgll3</sub>EE<sub>akap11</sub>*, ~25% *EL<sub>vgll3</sub>EL<sub>akap11</sub>*,<br>~25% *LL<sub>vgll3</sub>EL<sub>akap11</sub>*, ~25% *LL<sub>vgll3</sub>LL<sub>akap11</sub>* |
| *LL<sub>vgll3</sub>EL<sub>akap11</sub>* × *EL<sub>vgll3</sub>EE<sub>akap11</sub>* | 3 | ~25% *EL<sub>vgll3</sub>EE<sub>akap11</sub>*, ~25% *LL<sub>vgll3</sub>EE<sub>akap11</sub>*,<br>~25% *EL<sub>vgll3</sub>EL<sub>akap11</sub>*, ~25% *LL<sub>vgll3</sub>EL<sub>akap11</sub>* |
| *LL<sub>vgll3</sub>LL<sub>akap11</sub>* × *LL<sub>vgll3</sub>EL<sub>akap11</sub>* | 2 | ~50% *LL<sub>vgll3</sub>EL<sub>akap11</sub>*, ~50% *LL<sub>vgll3</sub>LL<sub>akap11</sub>* |
| *LL<sub>vgll3</sub>EE<sub>akap11</sub>* × *EL<sub>vgll3</sub>EE<sub>akap11</sub>* | 1 | ~50% *EL<sub>vgll3</sub>EE<sub>akap11</sub>*, ~50% *LL<sub>vgll3</sub>EE<sub>akap11</sub>* |
| *EL<sub>vgll3</sub>EE<sub>akap11</sub>* × *EE<sub>vgll3</sub>EE<sub>akap11</sub>* | 1 | ~50% *EE<sub>vgll3</sub>EE<sub>akap11</sub>*, ~50% *EL<sub>vgll3</sub>EE<sub>akap11</sub>* |

### Fish husbandry

Eggs were fertilized in November 2019 and incubated in mesh-separated compartments (to keep families separate) in vertical incubators with re-circulated water at a mean water temperature of 7.1°C. Compartments were randomly organized in the incubator. At the eyed-egg stage, each family was transferred to one of sixteen 285L tanks equipped with two water recirculation systems that have controlled water temperature, oxygen, and light conditions. Water parameters such as pH, ammonia, nitrite and nitrate were also monitored. Tank water temperature ranged from 5.2°C to 17.6°C (Supplementary Figure 1). Tank lighting followed the natural cycle that would occur at 62°24’N and 25°57’E. Fish were fed live Artemia for ten days and then fed commercial aquaculture feed ad libitum (Raisio Baltic Blend) for the remainder of the experiment. Size of feed pellets increased over time according to fish size. In 12 of the 16 tanks (those with the largest family sizes) 12mm passive integrated transponder tags were inserted into the body cavity, and a fin clip taken, during June-July 2019 following anaesthesia with methanesulfonate to enable re-identification and genotyping. Water temperature was decreased to 13°C for this period to reduce stress of fish due to handling. In order to keep the biomass of these 12 tanks at an acceptable level towards the end of the experiment, females were identified based on genotypic sex and culled July to September 2020. This strategy was chosen as only male Atlantic salmon are able to mature at one year of age in captivity (Debes et al. 2019) and therefore maximizing male numbers also maximizes sample sizes for the maturation phenotype. Nevertheless, a minimum of 40 females were retained in each tank. In some cases biomass levels became too high even following culling of females and therefore some males were randomly culled between September and November 2020.

### DNA Extraction & Genotyping

Fin clips from all individuals were placed directly into *Lucigen QuickExtract DNA Extraction Solution 1*.*0* to extract DNA. The *vgll3, akap11* and *SDY* loci were genotyped using the Kompetitive allele-specific polymerase chain reaction (KASP^*TM*^) method (He, Holme, and Anthony 2014). Two alternative allele specific forward primers and one reverse primer were designed by *LGC Biosearch Technologies* for the *vgll3* and *akap11* loci. An amplification/non-amplification assay was designed for the male specific SDY locus, and this assay also included primers for amplification of a region of the 18S locus as a positive control for assay performance (Supplementary Table 2). The reaction mix for each reaction consisted of 2.5 µl of sample DNA, 2.5 µl KASP 2x Master mix, 0.07 µl KASP Assay mix which contains the locus-specific primers. The reactions were performed with qPCR machines (C1000 Thermal cycler with CFX384 Real-Time System, Bio-Rad) and the following thermal cycling conditions: 94°C for 15 minutes (1 cycle); 94°C for 20 seconds, 61°C for 1 minutes and decreasing temperature by 0.6°C per cycle (10 cycles); 94°C for 20 seconds, 55°C for 1 minute (29 cycles); 37°C for 1 minute; 94°C for 20 seconds, 57°C for 1 minute (3 cycles); 37°C for 1 minute, read plate; and 4°C for 3 minutes. Genotypes of the *vgll3* and *akap11* SNPs were called using allelic discrimination implemented in the CFX Maestro software (*Bio-Rad*). Genotypic sex was determined by analyzing the per-individual difference between ROX-standardized FAM and HEX florescence values using the *normalmixEM* function in *mixtools R* package. Florescence of the FAM alleles indicates the presence of the SDY locus (Supplementary Table 2), which is male-specific in Atlantic salmon (Yano et al. 2012). An individual with a FAM-HEX value within two standard deviations from the mean of the upper normal distribution was considered a male. In contrast, individuals with a FAM-HEX value within two standard deviations of the lower normal distribution mean were considered female.

### Data collection

At the completion of the experiment during November and December, 2020, we recorded length (fork length), mass and maturity status (immature/mature) for all male individuals. To identify males, individuals were dissected and checked internally for the presence of male or female gonads. Maturity status was determined via examination of the gonads size and colour. Individuals were considered mature if the gonads were a milky white colour and enlarged so that they filled at least 75% of the body cavity. For a subset of individuals (N=632) that were kept alive for a different experiment and could not be dissected, we relied on genotypic sex. Maturity status for these males was determined by pressing on the abdomen and checking for the release of milt, which would indicate the male was mature.

### Data analysis

We tested for an association between maturation status in male Atlantic salmon and the genotypes of two adjacent genes, *vgll3* and *akap11*. Maturation status was modelled as a binary trait (immature=0, mature=1) using mixed-effect logistic regression implemented in *lme4 R* package. We first identified the most parsimonious null model, with no genetic terms, to fit the data. Fork length, Fulton’s condition factor and their interaction were included as fixed effects and family was included as a random effect. Fork length and Fulton’s condition factor were mean-centred. Using the *dredge* function in the *MuMin* package in *R* (Barton 2020), the most parsimonious model was selected based on each models corrected Akaike Information Criterion (AICc) scores. Genetic terms for the focal loci, *vgll3* and *akap11*, are then added to the selected model to test for an effect of these loci on maturation odds. We first modelled the effect of both genes on maturation status by including each locus as its own genetic term. The genetic terms were included as a categorical effect, rather than numerical, in order to not assume an additive genetic effect. We then examined the effect of combined genotypes on maturations odds by including genotypes at each gene as a single term in the model. We compared combined genotypes where alleles at one gene were the same and alleles at the other gene varied. Two models included genotypes where *akap11* genotype remained consistent but *vgll3* genotype varied: 1) *EE*_*vgll3*_*EE*_*akap11*_, *EL*_*vgll3*_*EE*_*akap11*_, *LL*_*vgll3*_*EE*_*akap11*_ and 2) *EL*_*vgll3*_*EL*_*akap11*_, *LL*_*vgll3*_*EL*_*akap11*_. The other two models included genotypes where *vgll3* genotype remained consistent but *akap11* genotype varied: 3) *EL*_*vgll3*_*EE*_*akap11*_, *EL*_*vgll3*_*EL*_*akap11*_ and 4) *LL*_*vgll3*_*EE*_*akap11*_, *LL*_*vgll3*_*EL*_*akap11*_, *LL*_*vgll3*_*LL*_*akap11*_.

## RESULTS

A total of 5895 males were raised until the end of the experiment. The overall maturation rate was 2.87%. Average mass, length and maturation rate of each family is listed in Supplementary Table 1. Of these 5895 individuals, 4769 had recombinant genotypes (i.e. carrying a haplotype with an *L* allele for *vgll3* and an *E* allele for *akap11*, or vice versa). The *E* allele frequencies of *vgll3* and *akap11* were 0.30 and 0.69, respectively.

The most parsimonious model explaining maturation status included length as a fixed effect and family as a random effect. *Vgll3* had a much stronger effect than *akap11* on maturation status, where the *vgll3 EE* and *EL* genotypes increased the *log*(odds ratio) of maturing relative to the *LL* genotype by 4.21 and 1.79, respectively. Contrastingly, only the *akap11 EE* genotype had a marginally significant negative effect on the odds of maturation, whereby it decreased the *log*(odds ratio) of maturing by 1.30 (Figure 1, Supplementary Table 3). Similarly, the effects of combined genotypes on the odds of maturation suggested a strong effect of all *vgll3* genotypes and a weak effect of the *akap11 EE* genotype. Allele changes at *vgll3* alter the odds of maturation for all observed genotype combinations, whereby genotypes with *vgll3 E* alleles increased the odds of maturation relative to those with the *L* allele (Figure 2a, b). In contrast allele changes at *akap11* altered the odds of maturation for only one genotype combination (*EL*_*vgll3*_*EE*_*akap11*_), whereby the *EL*_*vgll3*_*EE*_*akap11*_ genotype slightly decreased the odds of maturation relative to *EL*_*vgll3*_*EL*_*akap11*_ genotype (Figure 2c, d, Supplementary Tables 4-7).

**Figure 1.**
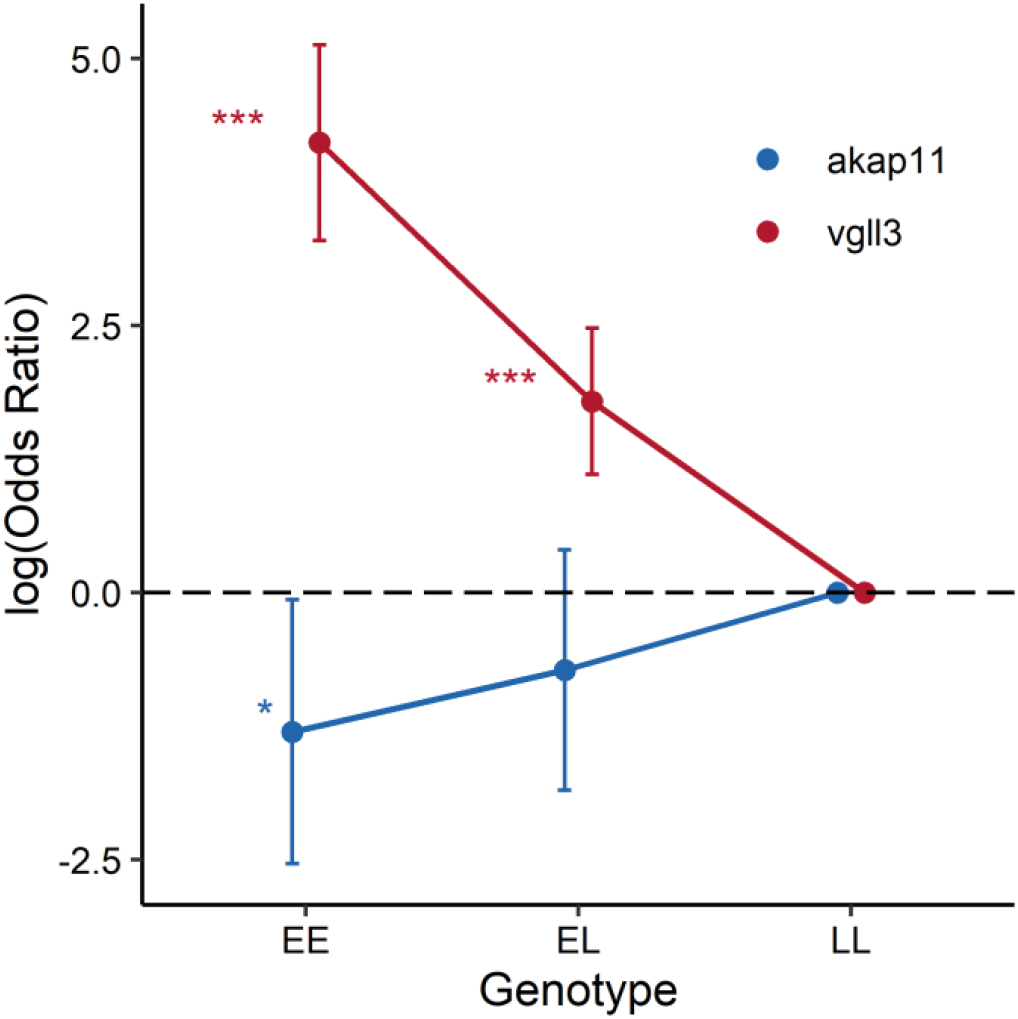
Ratio of the odds of maturation on the logarithmic scale and the respective 95% confidence intervals of the *EE* and *EL* genotypes for *vgll3* and *akap11*, relative to the *LL* genotype. Asterisks denote level of significance (* *p*-value < 0.05, *** *p*-value < 0.001). The *E* and *L* refer to the alleles associated with earlier and later maturation, respectively, in Barson et al (2015).

**Figure 2.**
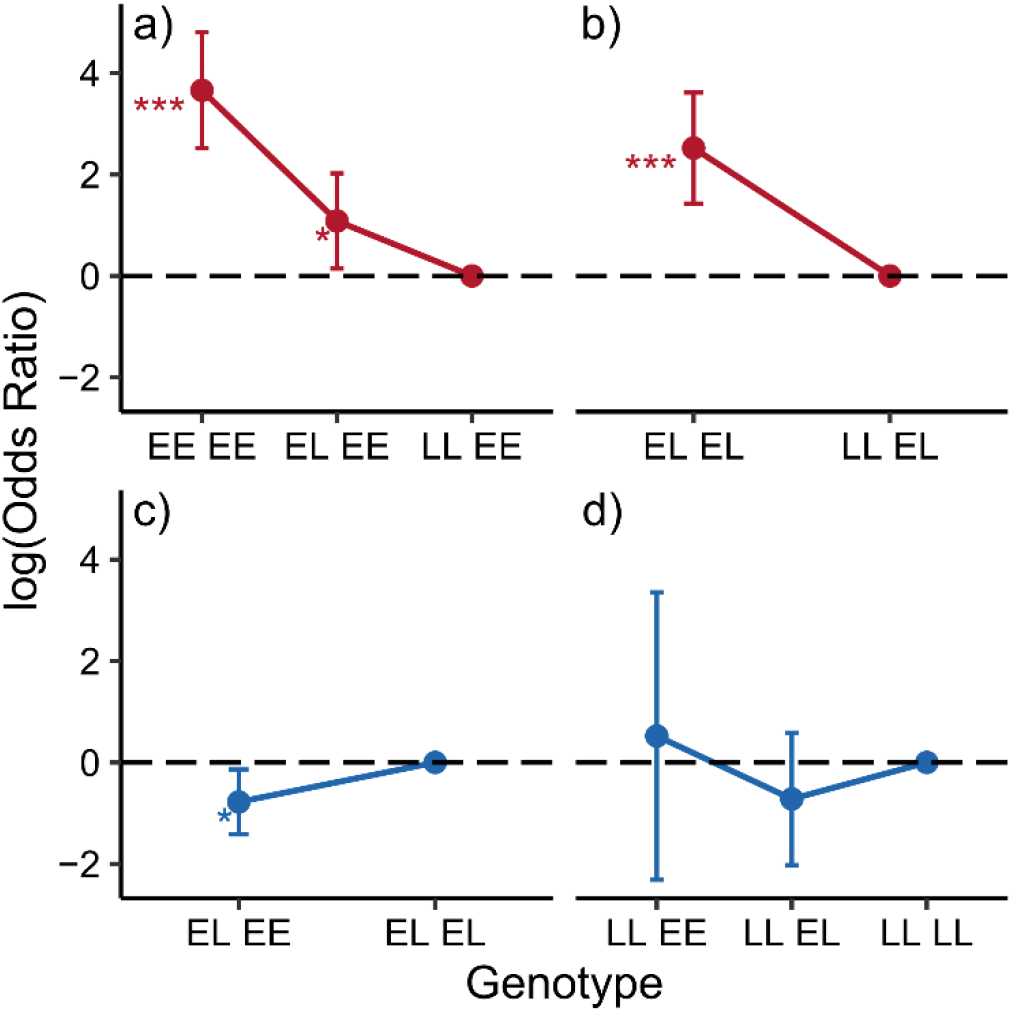
Ratio of the odds of maturation on the logarithmic scale and the respective 95% confidence intervals of the combined genotypes. Each comparison is plotted separately: a) *EE*_*vgll3*_*EE*_*akap11*_, *EL*_*vgll3*_*EE*_*akap11*_, *LL*_*vgll3*_*EE*_*akap11*_; b) *EL*_*vgll3*_*EL*_*akap11*_, *LL*_*vgll3*_*EL*_*akap11*_; c) *EL*_*vgll3*_*EE*_*akap11*_, *EL*_*vgll3*_*EL*_*akap11*_; and d) *LL*_*vgll3*_*EE*_*akap11*_, *LL*_*vgll3*_*EL*_*akap11*_, *LL*_*vgll3*_*LL*_*akap11*_. Estimates within each comparison are relative to the genotype with the most *L* alleles. Asterisks denote level of significance (* *p*-value < 0.05, *** *p*-value < 0.001). The first and last two alleles listed indicate the *vgll3* and *akap11* genotypes, respectively, where *E* and *L* are the alleles found to be associated with earlier and later maturation, respectively, in Barson et al. (2015).

## DISCUSSION

Previous genome-wide association studies (GWAS), found a strong association between maturation timing in Atlantic salmon and a region on chromosome 25. These studies have shown that significantly associated SNPs span a ∼250kb genomic with the strongest association signal occurring between two genes, *vgll3* and *akap11*. Due to linkage disequilibrium within the region, it remained unclear which SNPs were potentially causal and which were spuriously associated via linkage. Here, we took advantage of existing recombination events to breed a large set of progeny with reduced linkage disequilibrium between potential causal SNPs at the candidate locus. We found that SNP variation within *akap11* has little to no effect on maturation timing and therefore the effect of the locus is primarily driven by variation in closer proximity to *vgll3*. This refines the genomic location of SNP variation affecting Atlantic salmon maturation timing at a key large-effect locus and improves our understanding of the gene variation most likely underlying differences in maturation timing. These findings will help guide future experimentation determining the role of this large-effect locus in genetic processes involved in Atlantic salmon maturation.

Here, we measured the effect of *vgll3* based on the genotype of a SNP 7.5kb downstream of *vgll3* that showed the strongest association in 57 wild Atlantic salmon populations (Barson et al. 2015). This genotype was in complete linkage disequilibrium with the genotypes of the two missense mutations within *vgll3* due to our parent selection criteria. Regardless of the accompanying *akap11* genotype, the *vgll3* genotype had a strong effect on maturation timing, with the *E* allele showing similar strong positive effects on early maturation. In contrast, *akap11* genotype showed a relatively small effect on maturation where maturation odds unexpectedly decreased in *EL* and *EE* individuals relative to *LL* individuals. If the effect of *akap11* variation is true, its relative contribution to controlling maturation timing would be minimal given the effect is both small and found in only one genotype class. These results provide convincing evidence that variation closer to *vgll3* than *akap11* is linked with maturation timing, at least in male parr. However, it is important to recognize that we do not yet know if any of the *vgll3*-SNPs are causal themselves, or simply linked to causal variation. Additionally, we cannot rule out the possibility that the causal variation driving the *vgll3* genotype effect alters the functioning of a different gene, however given the tight genetic linkage of the *vgll3*-SNPs and *vgll3, vgll3* variation is nevertheless strongly associated with maturation timing. Recent functional research also supports this notion - *vgll3* expression in immature testes of Atlantic salmon differs between *vgll3* genotypes, *EE* and *LL* (Verta et al. 2020), which suggests that SNP variation linked with *vgll3* also associates with altered *vgll3* function. Further, we cannot exclude the possibility that the causal variation may regulate both *vgll3* and *akap11* given their close proximity. Shared regulatory regions are prevalent in the human genome (Trinklein et al. 2004). Interestingly, expression of *vgll3* and *akap11* are correlated during early development (Kurko et al. 2020). Examining genotype-specific expression levels of *akap11* and *vgll3* in recombinant individuals may help to further resolve the functional significance of the causal variation at this large-effect locus.

No recombination events introducing haplotypes with the *vgll3 E* and *akap11 L* alleles were found in the parental source. Therefore, there were no progeny with genotypes *EE*_*vgll3*_*EL* and *EE*_*vgll3*_*LL* and we were thus unable to test the effect of these genotype combinations. Furthermore, it is unclear whether the haplotypes with *vgll3 L* and *akap11 E* alleles found here arose via a single recombination event, or multiple events. Sequencing of this region in the parental individuals would identify the location of recombination breakpoint(s) and therefore the number of recombination events. The location of breakpoint(s) also helps to narrow down the causal region, as any variation downstream of the breakpoint can be ruled out.

A further caveat of our study is that due to large number of individuals raised, we did not have resources for genotyping a sufficient number of loci for parentage assignment and were thus unable to randomize individuals across tanks. For this reason, we are unable to tease apart tank effects and family effects and therefore the effect of the term “family” in our models also includes any tank effects. To help account for this we included multiple families for most of the cross types, which ensured each of the genotype combinations were raised in multiple tanks. We also expect that family effects would account for a substantial portion of the variation explained by the family/tank model term given the polygenic architecture of Atlantic salmon maturation (Sinclair-Waters et al. 2020). Additionally, Debes et al. (2019) randomized individuals from many families across multiple tanks and found inter-family variation in maturation rate and no effect of tank.

Our findings suggest that *vgll3* would be an appropriate target for knockout with CRISPR to further resolve the effect of *vgll3* on Atlantic salmon maturation. Genome editing with CRISPR-*Cas9* has successfully generated Atlantic salmon with gene knock-outs (Wargelius et al. 2016; Edvardsen et al. 2014). Further, in other species, the variants causing trait variation have been finely mapped to a single or set of mutation(s) with CRISPR-*Cas9* genome editing (Karageorgi et al. 2019; Li et al. 2020; Ward et al. 2021). Given the large-effect of the *vgll3* locus, it would be an interesting focus for fine-mapping with genome editing technology, whereby effects of the two missense mutations and the top-associated SNPs from previous association studies could be delineated. Single base editing of the known missense mutation and top-associated SNPs could introduce novel genotype combinations and help to more finely map and/or validate causal mutations at the *vgll3* locus. Some success with single base editing in Atlantic salmon has been accomplished, whereby 30% to 60% efficiency was achieved (Straume et al. 2021), suggesting editing of *vgll3* SNPs may be feasible. Alternatively, identifying individuals carrying natural recombinant alleles at the *vgll3* locus may be possible, however this may require genotyping and scanning a large number individuals from many source populations, followed by rearing of males and females to maturation age.

As genome assembly and genomic data production for species in the wild becomes easier, the number of candidate loci linked to association and/selection signals is likely to rise. Our findings demonstrate how, when identified, natural recombinants can be used to more precisely map causal variation underlying such signals when phenotypic data can be obtained. Furthermore, offspring, from controlled crosses maximizing the number of recombinants, could potentially be released into the wild, which would allow for follow-up studies in the wild. Such follow-up studies in the wild are unlikely possible if CRISPR or other genome editing technology is used. Another approach that takes advantage of natural recombination events, admixture mapping (Vasemägi and Primmer 2005; Mckeigue 1998), can be applied in a natural setting and is thus a promising method in systems where raising individuals in captivity is not feasible. Admixture mapping, however, relies on hybridization between populations with different allele frequencies at trait-associated loci and therefore can only be applied under these specific conditions. In conclusion, using natural recombination events to narrow down the genomic location of causal variation of ecologically relevant traits is in effective approach and can be especially useful in systems where genome editing is particular challenging or not feasible.

## Acknowledgements

Funding was provided by Academy of Finland (grant numbers 307593, 302873 and 327255), the European Research Council under the European Articles Union’s Horizon 2020 research and innovation program (grant no. 742312) and a Natural Sciences and Engineering Research Council of Canada postgraduate scholarship. We would like to acknowledge staff at the Natural Resources Institute Finland hatchery in Laukaa and members of the Evolution, Conservation and Genomics research group for their help in coordinating and collecting gametes for crosses. We acknowledge Shadi Jansouz for performing DNA extractions. We thank Paul Debes for sharing advice and experience from prior experiments on salmon maturation. We thank Nea Asumaa, Eirini Karathanou, Clelia Mulà, and Seija Tillanen for help with fish husbandry. We also thank Nea Asumaa, Iikki Donner, Dorian Jagusch, Eirini Karathanou, Jacqueline Moustakas-Verho, Clelia Mulà, Outi Ovaskainen, Seija Tillanen, and Valeria Valanne for help with tagging, tissue sampling, and phenotyping.

## Data Accessibility

Genotypes and phenotypic data that support the findings of this study are openly available in Dryad at http://doi.org/[doi].

## Authors’ Contributions

CRP, MSW conceived the study. MSW, CRP designed crosses. CRP, NP, MSW designed experimental setup. NP supervised fish husbandry and maintenance of fish-raising facility. MSW, NP led tagging, tissue collection, and phenotypic data collection. AR, TA, CRP developed the KASP genotyping protocol. AR performed genotyping. MSW performed genotype calling and data analysis. JE provided parental material for crosses. MSW, CRP drafted the manuscript. All authors approved the final version of the manuscript.

## References

Albadri, Shahad, Flavia De Santis, Vincenzo Di Donato, and Filippo Del Bene. 2017. “CRISPR/Cas9-Mediated Knockin and Knockout in Zebrafish.” In, edited by Rudolf Jaenisch, Feng Zhang, and Fred Gage, 41–49. Cham (CH). https://doi.org/10.1007/978-3-319-60192-2_4.

Aykanat, T., M. Lindqvist, V. L. Pritchard, and C. R. Primmer. 2016. “From Population Genomics to Conservation and Management: A Workflow for Targeted Analysis of Markers Identified Using Genome-Wide Approaches in Atlantic Salmon Salmo Salar.” Journal of Fish Biology, no. 89: 2658–79. https://doi.org/10.1111/jfb.13149.

Ayllon, Fernando, Erik Kjærner-Semb, Tomasz Furmanek, Vidar Wennevik, Monica F. Solberg, Geir Dahle, Geir Lasse Taranger, et al. 2015. “The Vgll3 Locus Controls Age at Maturity in Wild and Domesticated Atlantic Salmon (Salmo Salar L.) Males.” PLoS Genetics 11 (11): 1–15. https://doi.org/10.1371/journal.pgen.1005628.

Barson, Nicola J, Tutku Aykanat, Kjetil Hindar, Matthew Baranski, Geir H Bolstad, Peder Fiske, Céleste Jacq, et al. 2015. “Sex-Dependent Dominance at a Single Locus Maintains Variation in Age at Maturity in Salmon.” Nature 528 (7582): 405–8. https://doi.org/10.1038/nature16062.

Barton, K. 2020. “MuMIn: Multi-Model Inference. R Package Version 1.43.17.”

Concha, Carolina, Richard W.R. Wallbank, Joseph J. Hanly, Jennifer Fenner, Luca Livraghi, Edgardo Santiago Rivera, Daniel F. Paulo, et al. 2019. “Interplay between Developmental Flexibility and Determinism in the Evolution of Mimetic Heliconius Wing Patterns.” Current Biology 29 (23): 3996-4009.e4. https://doi.org/10.1016/j.cub.2019.10.010.

Cousminer, Diana L., Diane J. Berry, Nicholas J. Timpson, Wei Ang, Elisabeth Thiering, Enda M. Byrne, H. Rob Taal, et al. 2013. “Genome-Wide Association and Longitudinal Analyses Reveal Genetic Loci Linking Pubertal Height Growth, Pubertal Timing and Childhood Adiposity.” Human Molecular Genetics 22 (13): 2735–47. https://doi.org/10.1093/hmg/ddt104.

Day, Felix R., Deborah J. Thompson, Hannes Helgason, Daniel I. Chasman, Hilary Finucane, Patrick Sulem, Katherine S. Ruth, et al. 2017. “Genomic Analyses Identify Hundreds of Variants Associated with Age at Menarche and Support a Role for Puberty Timing in Cancer Risk.” Nature Genetics 49 (6): 834–41. https://doi.org/10.1038/ng.3841.

Debes, Paul V, Nikolai Piavchenko, Annukka Ruokolainen, Jacqueline E Moustakas-verho, Noora Parre, Tutku Aykanat, Jaakko Erkinaro, Craig R Primmer, and Paul V Debes. 2019. “Large Single-Locus Effects for Maturation Timing Are Mediated via Body Condition in Atlantic Salmon.” BioRxiv. https://doi.org/10.1101/780437.

Edvardsen, Rolf B., Sven Leininger, Lene Kleppe, Kai Ove Skaftnesmo, and Anna Wargelius. 2014. “Targeted Mutagenesis in Atlantic Salmon (Salmo Salar L.) Using the CRISPR/Cas9 System Induces Complete Knockout Individuals in the F0 Generation.” PLoS ONE 9 (9). https://doi.org/10.1371/journal.pone.0108622.

Erkinaro, Jaakko, Yann Czorlich, Panu Orell, Jorma Kuusela, Morten Falkegård, Maija Länsman, Henni Pulkkinen, Craig R Primmer, and Eero Niemelä. 2019. “Life History Variation across Four Decades in a Diverse Population Complex of Atlantic Salmon in a Large Subarctic River.” Canadian Journal of Fisheries and Aquatic Sciences 76 (1): 42–55. https://doi.org/10.1139/cjfas-2017-0343.

Fleming, I. A., and S Einum. 2011. “Reproductive Ecology: A Tale of Two Sexes.” In Atlantic Salmon Ecology, 35–65.

Good, Christopher, and John Davidson. 2016. “A Review of Factors Influencing Maturation of Atlantic Salmon, Salmo Salar, with Focus on Water Recirculation Aquaculture System Environments.” Journal of the World Aquaculture Society 47 (5): 605–32. https://doi.org/10.1111/jwas.12342.

Halperin, Daniel S., Calvin Pan, Aldons J. Lusis, and Peter Tontonoz. 2013. “Vestigial-like 3 Is an Inhibitor of Adipocyte Differentiation.” Journal of Lipid Research 54 (2): 473–81. https://doi.org/10.1194/jlr.m032755.

He, Chunlin, John Holme, and Jeffrey Anthony. 2014. “SNP Genotyping: The KASP Assay BT - Crop Breeding: Methods and Protocols.” In, edited by Delphine Fleury and Ryan Whitford, 75–86. New York, NY: Springer New York. https://doi.org/10.1007/978-1-4939-0446-4_7.

Johnston, Susan E., John C. McEwan, Natalie K. Pickering, James W. Kijas, Dario Beraldi, Jill G. Pilkington, Josephine M. Pemberton, and Jon Slate. 2011. “Genome-Wide Association Mapping Identifies the Genetic Basis of Discrete and Quantitative Variation in Sexual Weaponry in a Wild Sheep Population.” Molecular Ecology 20 (12): 2555–66. https://doi.org/10.1111/j.1365-294X.2011.05076.x.

Johnston, Susan E., Panu Orell, Victoria L. Pritchard, Matthew P. Kent, Sigbjørn Lien, Eero Niemelä, Jaakko Erkinaro, and Craig R. Primmer. 2014. “Genome-Wide SNP Analysis Reveals a Genetic Basis for Sea-Age Variation in a Wild Population of Atlantic Salmon (Salmo Salar).” Molecular Ecology 23 (14): 3452–68. https://doi.org/10.1111/mec.12832.

Karageorgi, Marianthi, Simon C Groen, Fidan Sumbul, Julianne N Pelaez, Kirsten I Verster, Jessica M Aguilar, Amy P Hastings, et al. 2019. “Genome Editing Retraces the Evolution of Toxin Resistance in the Monarch Butterfly.” Nature. https://doi.org/10.1038/s41586-019-1610-8.

Kardos, Marty, Gordon Luikart, Rowan Bunch, Sarah Dewey, William Edwards, Sean McWilliam, John Stephenson, Fred W. Allendorf, John T. Hogg, and James Kijas. 2015. “Whole-Genome Resequencing Uncovers Molecular Signatures of Natural and Sexual Selection in Wild Bighorn Sheep.” Molecular Ecology 24 (22): 5616–32. https://doi.org/10.1111/mec.13415.

Kurko, Johanna, Paul V. Debes, Andrew H. House, Tutku Aykanat, Jaakko Erkinaro, and Craig R. Primmer. 2020. “Transcription Profiles of Age-at-Maturity-Associated Genes Suggest Cell Fate Commitment Regulation as a Key Factor in the Atlantic Salmon Maturation Process.” G3 (Bethesda, Md.) 10 (1): 235–46. https://doi.org/10.1534/g3.119.400882.

Li, Feng, Akira Komatsu, Miki Ohtake, Heesoo Eun, Akemi Shimizu, and Hiroshi Kato. 2020. “Direct Identification of a Mutation in OsSh1 Causing Non-Shattering in a Rice (Oryza Sativa L.) Mutant Cultivar Using Whole-Genome Resequencing.” Scientific Reports 10 (1): 1–13. https://doi.org/10.1038/s41598-020-71972-1.

Livraghi, Luca, Arnaud Martin, Melanie Gibbs, Nora Braak, Saad Arif, and Casper J. Breuker. 2018. CRISPR/Cas9 as the Key to Unlocking the Secrets of Butterfly Wing Pattern Development and Its Evolution. Advances in Insect Physiology. 1st ed. Vol. 54. Elsevier Ltd. https://doi.org/10.1016/bs.aiip.2017.11.001.

Luconi, Michaela, Vinicio Carloni, Fabio Marra, Pietro Ferruzzi, Gianni Forti, and Elisabetta Baldi. 2004. “Increased Phosphorylation of AKAP by Inhibition of Phosphatidylinositol 3-Kinase Enhances Human Sperm Motility through Tail Recruitment of Protein Kinase A.” Journal of Cell Science 117 (Pt 7): 1235–46. https://doi.org/10.1242/jcs.00931.

Mckeigue, Paul M. 1998. “Mapping Genes That Underlie Ethnic Differences in Disease Risk : Methods for Detecting Linkage in Admixed Populations, by Conditioning on Parental Admixture,” 241–51.

Miki, Kiyoshi, William D Willis, Paula R Brown, Eugenia H Goulding, Kerry D Fulcher, and Edward M Eddy. 2002. “Targeted Disruption of the Akap4 Gene Causes Defects in Sperm Flagellum and Motility.” Developmental Biology 248 (2): 331–42. https://doi.org/10.1006/dbio.2002.0728.

Mobley, K.B., Hanna Granroth-Wilding, Mikko Ellmén, Panu Orell, Jaakko Erkinaro, and Craig R. Primmer. 2020. “Time Spent in Distinct Life History Stages Has Sex-Specific Effects on Reproductive Fitness in Wild Atlantic Salmon.” Molecular Ecology 29 (6): 1173–84. https://doi.org/10.1111/mec.15390.

Mobley, Kenyon B., Tutku Aykanat, Yann Czorlich, Andrew House, Johanna Kurko, Antti Miettinen, Jacqueline Moustakas-Verho, et al. 2021. Maturation in Atlantic Salmon (Salmo Salar, Salmonidae): A Synthesis of Ecological, Genetic, and Molecular Processes. Reviews in Fish Biology and Fisheries. Vol. 0123456789. https://doi.org/10.1007/s11160-021-09656-w.

Perry, John R B, Felix Day, Cathy E Elks, Patrick Sulem, Deborah J Thompson, Teresa Ferreira, Chunyan He, et al. 2014. “Parent-of-Origin-Specific Allelic Associations among 106 Genomic Loci for Age at Menarche.” Nature 514 (July): 92. https://doi.org/http://10.0.4.14/nature13545.

Pritchard, V. L., Makinen H., Vähä J.-P., Erkinaro J., Orell P., and C.R. Primmer. 2018. “Genomic Signatures of Fine-Scale Selection in Atlantic Salmon Suggest Involvment of Sexual Maturation, Energy Homeostatis, and Immune Defence-Related Genes.” Molecular Ecology 27: 2560–75. https://doi.org/10.1111/mec.14705.

Reinton, Nils, Philippe Collas, Trine B Haugen, Bjørn S Skålhegg, Vidar Hansson, Tore Jahnsen, and Kjetil Taskén. 2000. “Localization of a Novel Human A-Kinase-Anchoring Protein, HAKAP220, during Spermatogenesis.” Developmental Biology 223 (1): 194–204. https://doi.org/10.1006/dbio.2000.9725.

Rodríguez-Leal, Daniel, Zachary H Lemmon, Jarrett Man, Madelaine E Bartlett, and Zachary B Lippman. 2017. “Engineering Quantitative Trait Variation for Crop Improvement by Genome Editing.” Cell 171 (2): 470-480.e8. https://doi.org/10.1016/j.cell.2017.08.030.

Sander, Jeffry D, and J Keith Joung. 2014. “CRISPR-Cas Systems for Editing, Regulating and Targeting Genomes.” Nature Biotechnology 32 (4): 347–55. https://doi.org/10.1038/nbt.2842.

Sedeek, Khalid E M, Ahmed Mahas, and Magdy Mahfouz. 2019. “Plant Genome Engineering for Targeted Improvement of Crop Traits.” Frontiers in Plant Science 10 (February): 114. https://doi.org/10.3389/fpls.2019.00114.

Sinclair-Waters, Marion, Ian R. Bradbury, Corey J. Morris, Sigbjørn Lien, Matthew P. Kent, and Paul Bentzen. 2017. “Ancient Chromosomal Rearrangement Associated with Local Adaptation of a Post-Glacially Colonized Population of Atlantic Cod in the Northwest Atlantic.” Molecular Ecology, no. October: 1–13. https://doi.org/10.1111/mec.14442.

Sinclair-Waters, Marion, Torfinn Nome, Jing Wang, Sigbjørn Lien, Matthew P Kent, Harald Sægrov, Bjørn Florø-Larsen, Geir H Bolstad, Craig R Primmer, and Nicola J Barson. 2021. “Dissecting the Loci Underlying Maturation Timing in Atlantic Salmon Using Haplotype and Multi-SNP Based Association Methods.” BioRxiv. https://doi.org/10.1101/2021.05.28.446127.

Sinclair-Waters, Marion, Jørgen Ødegård, Sven Arild Korsvoll, Thomas Moen, Sigbjørn Lien, Craig R. Primmer, and Nicola J. Barson. 2020. “Beyond Large-Effect Loci: Large-Scale GWAS Reveals a Mixed Large-Effect and Polygenic Architecture for Age at Maturity of Atlantic Salmon.” Genetics, Selection, Evolution : GSE 52 (1): 9. https://doi.org/10.1186/s12711-020-0529-8.

Straume, Anne Hege, Erik Kjærner-Semb, Kai Ove Skaftnesmo, Hilal Güralp, Simon Lillico, Anna Wargelius, and Rolf Brudvik Edvardsen. 2021. “A Refinement to Gene Editing in Atlantic Salmon Using Asymmetrical Oligonucleotide Donors.” bioRxiv. https://doi.org/10.1101/2021.02.08.430296.

The International Mouse Knockout Consortium. 2007. “A Mouse for All Reasons.” Cell 128 (1): 9–13. https://doi.org/10.1016/j.cell.2006.12.018.

Trinklein, Nathan D., Shelley Force Aldred, Sara J. Hartman, Diane I. Schroeder, Robert P. Otillar, and Richard M. Myers. 2004. “An Abundance of Bidirectional Promoters in the Human Genome.” Genome Research 14 (1): 62–66. https://doi.org/10.1101/gr.1982804.

Varshney, Gaurav K, Jing Lu, Derek E Gildea, Haigen Huang, Wuhong Pei, Zhongan Yang, Sunny C Huang, et al. 2013. “A Large-Scale Zebrafish Gene Knockout Resource for the Genome-Wide Study of Gene Function.” Genome Research 23 (4): 727–35. https://doi.org/10.1101/gr.151464.112.

Vasemägi, A., and C. R. Primmer. 2005. “Challenges for Identifying Functionally Important Genetic Variation: The Promise of Combining Complementary Research Strategies.” Molecular Ecology 14 (12): 3623–42. https://doi.org/10.1111/j.1365-294X.2005.02690.x.

Verta, Jukka-Pekka, Paul Vincent Debes, Nikolai Piavchenko, Annukka Ruokolainen, Outi Ovaskainen, Jacqueline Emmanuel Moustakas-Verho, Seija Tillanen, et al. 2020. “Cis-Regulatory Differences in Isoform Expression Associate with Life History Strategy Variation in Atlantic Salmon.” PLOS Genetics 16 (9): e1009055. https://doi.org/10.1371/journal.pgen.1009055.

Ward, Christopher M., Roswitha A. Aumann, Mark A. Whitehead, Katerina Nikolouli, Gary Leveque, Georgia Gouvi, Elisabeth Fung, et al. 2021. “White Pupae Phenotype of Tephritids Is Caused by Parallel Mutations of a MFS Transporter.” Nature Communications 12 (1): 1–12. https://doi.org/10.1038/s41467-020-20680-5.

Wargelius, Anna, Sven Leininger, Kai Ove Skaftnesmo, Lene Kleppe, Eva Andersson, Geir Lasse Taranger, Rudiger W. Schulz, and Rolf B. Edvardsen. 2016. “Dnd Knockout Ablates Germ Cells and Demonstrates Germ Cell Independent Sex Differentiation in Atlantic Salmon.” Scientific Reports 6 (5817): 1–8. https://doi.org/10.1038/srep21284.

Woronik, Alyssa,Kalle Tunström, Michael W. Perry, Ramprasad Neethiraj, Constanti Stefanescu, Maria de la Paz Celorio-Mancera, Oskar Brattström, et al. 2019. “A Transposable Element Insertion Is Associated with an Alternative Life History Strategy.” Nature Communications 10 (1). https://doi.org/10.1038/s41467-019-13596-2.

Wucherpfennig, Julia I., Craig T. Miller, and David M. Kingsley. 2019. “Efficient CRISPR-Cas9 Editing of Major Evolutionary Loci in Sticklebacks.” Evolutionary Ecology Research 20 (1– 3): 1–29.

Yano, Ayaka, Barbara Nicol, Elodie Jouanno, Edwige Quillet, and Alexis Fostier. 2012. “The Sexually Dimorphic on the Y-Chromosome Gene (SdY) Is a Conserved Male-Specific Y-Chromosome Sequence in Many Salmonids.” Evolutionary Applications. https://doi.org/10.1111/eva.12032.

