## Supplementary Tables for "Refining the genomic location of SNP variation affecting Atlantic salmon maturation timing at a key large-effect locus"

Supplementary Table 1. Information for each family raised including sire and dam ID, genotype, their respective grandparent IDs, tank and water system. Data from final sampling of males including number raised, mean length, mean mass and maturation rate for each family.

| Family ID | Dam ID | Dam genotype | Dam's grandparents | Sire ID | Sire genotype | Sire grandparents | Tank | Water system | Number of males raised | Mean length (mm) | Mean mass (g) | Maturation rate |
|---|---|---|---|---|---|---|---|---|---|---|---|---|
| 1 | 3736 | $EL_{vgll3}EE_{akap11}$ | S09 & D06 | 3271 | $EL_{vgll3}EL_{akap11}$ | S11 & D01 | 7 | A | 383 | 134 | 30.4 | 0.211 |
| 2 | 3901 | $EL_{vgll3}EL_{akap11}$ | S08 & D13 | 3585 | $EL_{vgll3}EE_{akap11}$ | S02 & D02 | 12 | B | 372 | 139 | 32.7 | 0.056 |
| 3 | 3881 | $EL_{vgll3}EE_{akap11}$ | S01 & D09 | 3494 | $EL_{vgll3}EL_{akap11}$ | S06 & D12 | 13 | A | 190 | 120 | 22.3 | 0.032 |
| 4 | 3923 | $EL_{vgll3}EL_{akap11}$ | S06 & D01 | 3627 | $LL_{vgll3}EL_{akap11}$ | S02 & D12 | 5 | B | 354 | 138 | 33.0 | 0 |
| 5 | 3419 | $LL_{vgll3}EL_{akap11}$ | S01 & D13 | 3272 | $EL_{vgll3}EL_{akap11}$ | S05 & D04 | 1 | B | 367 | 130 | 26.9 | 0 |
| 6 | 3650 | $EL_{vgll3}EL_{akap11}$ | S11 & D14 | 3913 | $LL_{vgll3}EL_{akap11}$ | S04 & D11 | 2 | A | 440 | 132 | 28.4 | 0 |
| 7 | 3349 | $LL_{vgll3}EL_{akap11}$ | S04 & D11 | 3709 | $EL_{vgll3}EL_{akap11}$ | S02 & D03 | 6 | B | 342 | 141 | 34.8 | 0.003 |
| 8 | 3170 | $LL_{vgll3}EL_{akap11}$ | S08 & D01 | 3542 | $EL_{vgll3}EL_{akap11}$ | S10 & D02 | 9 | A | 265 | 136 | 32.2 | 0.011 |
| 9 | 3817 | $LL_{vgll3}EL_{akap11}$ | S08 & D13 | 3584 | $EL_{vgll3}EL_{akap11}$ | S06 & D12 | 4 | A | 512 | 136 | 30.3 | 0.002 |
| 10 | Unknown | $EE_{vgll3}EE_{akap11}$ or $EL_{vgll3}EE_{akap11}$ | Unknown | Unknown | $EE_{vgll3}EE_{akap11}$ or $EL_{vgll3}EE_{akap11}$ | Unknown | 3 | B | 295 | 133 | 29.9 | 0.041 |
| 11 | 3827 | $EL_{vgll3}EE_{akap11}$ | S09 & D11 | 3627 | $LL_{vgll3}EL_{akap11}$ | S02 & D12 | 10 | A | 362 | 146 | 37.3 | 0 |
| 12 | 3892 | $EL_{vgll3}EE_{akap11}$ | S08 & D09 | 3567 | $LL_{vgll3}EL_{akap11}$ | S04 & D14 | 8 | B | 468 | 126 | 22.7 | 0.004 |
| 13 | 3463 | $EL_{vgll3}EE_{akap11}$ | S02 & D09 | 3990 | $LL_{vgll3}EL_{akap11}$ | S08 & D13 | 11 | B | 406 | 141 | 36.2 | 0 |
| 14 | 3237 | $EL_{vgll3}EE_{akap11}$ | S04 & D13 | 3468 | $LL_{vgll3}EE_{akap11}$ | S01 & D11 | 15 | A | 353 | 135 | 28.7 | 0.057 |
| 15 | 3497 | $LL_{vgll3}EE_{akap11}$ | S08 & D06 | 3891 | $EL_{vgll3}EE_{akap11}$ | S02 & D01 | 14 | B | 572 | 136 | 30.4 | 0.007 |
| 16 | 3523 | $LL_{vgll3}LL_{akap11}$ | S03 & D02 | 3378 | $LL_{vgll3}EL_{akap11}$ | S02 & D13 | 16 | A | 214 | 129 | 26.3 | 0 |

Supplementary Table 2. Locus and primer information for genotyping assays used in this study. The target SNP alleles of the alternative forward primer sequences are indicated with square brackets. The fluorescence dye associated with the earlier and later maturation alleles are marked *E* and *L*, respectively. A region of 18S is amplified as a positive control in the amplification/non-amplification assay targeting SDY.

| Assayed genomic region | Assay type | Fluorescence dye | Forward primer | Reverse primer | Product size |
|---|---|---|---|---|---|
| *vgll3$_{TOP}$* | Allele specific PCR | Fam (L) | CCTCTGTTGTCATCCA GAATTAATC[A] | CACAGCTGTTCTGTACT GGAGGAAT | 57 bp |
| | | Hex (E) | CCTCTGTTGTCATCCA GAATTAATC[C] | | 57 bp |
| *akap11* | Allele specific PCR | Fam (L) | ATCTCCATGGAAACCAGCGTGAT[A] | TGATCCTGGAACCTGATAAATCCTTGTT | 84 bp |
| | | Hex (E) | CTCCATGGAAACCAGCGTGAT[G] | | 82 bp |
| SDY/18S | Amplification/non-amplification assay | Fam (target) | AGTACTGCGAAGAGGAGGTGCTTA | GAAGGGCTTGTAGGCAATTCTGACAT | 71 bp |
| | | Hex (control) | GTTGGTGGAGCGATTTGTCTGGTTA | GCTCGGGGCCGCATAACTAGTT | 78 bp |

Supplementary Table 3. Coefficients of the model examining the effects of both *vgll3* and *akap11* genotypes on the odds of maturation. Genotype estimates are relative to the *LL* genotype.

| Coefficient | Estimate | SE | z-value | p-value |
|---|---|---|---|---|
| Intercept | -6.28 | 0.77 | -8.18 | 2.79e-16 |
| *vgll3 EL* | 1.79 | 0.35 | 5.13 | 2.95e-07 |
| *vgll3 EE* | 4.21 | 0.47 | 9.01 | 2.06e-19 |
| *akap11 EL* | -0.72 | 0.57 | -1.26 | 2.07e-01 |
| *akap11 EE* | -1.30 | 0.63 | -2.06 | 3.91e-02 |
| Mean-centred length | -0.09 | 0.01 | -12.04 | 2.24e-33 |

Supplementary Table 4. Coefficients of the model comparing the effects of $EL_{vgll3}EE_{akap11}$ and $EL_{vgll3}El_{akap11}$ genotypes on the odds of maturation. Genotype estimate is relative to the $EL_{vgll3}EL_{akap11}$ genotype.

| Coefficient | Estimate | SE | z-value | p-value |
|---|---|---|---|---|
| Intercept | -5.06 | 0.68 | -7.42 | 1.22e-13 |
| $EL_{vgll3}EE_{akap11}$ | -0.78 | 0.33 | -2.38 | 1.73e-02 |
| Mean-centred length | -0.10 | 0.01 | -9.05 | 1.42e-19 |

Supplementary Table 5. Coefficients of the model comparing the effects of $LL_{vgll3}EE_{akap11}$, $LL_{vgll3}EL_{akap11}$ and $LL_{vgll3}LL_{akap11}$ genotypes on the odds of maturation. Genotype estimates are relative to the $LL_{vgll3}LL_{akap11}$ genotype.

| Coefficient | Estimate | SE | z-value | p-value |
|---|---|---|---|---|
| Intercept | -7.55 | 1.30 | -5.80 | 6.53e-09 |
| $LL_{vgll3}EL_{akap11}$ | -0.72 | 0.67 | -1.08 | 2.80e-01 |
| $LL_{vgll3}EE_{akap11}$ | 0.52 | 1.45 | 0.36 | 7.18e-01 |
| Mean-centred length | -0.09 | 0.01 | -6.13 | 8.69e-10 |

Supplementary Table 6. Coefficients of the model comparing the effects of $EE_{vgll3}EE_{akap11}$, $EL_{vgll3}EE_{akap11}$ and $LL_{vgll3}EE_{akap11}$ genotypes on the odds of maturation. Genotype estimates are relative to the $LL_{vgll3}EE_{akap11}$ genotype.

| Coefficient | Estimate | SE | z-value | p-value |
|---|---|---|---|---|
| Intercept | -6.53 | 0.77 | -8.56 | 1.17e-17 |
| $EL_{vgll3}EE_{akap11}$ | 1.08 | 0.48 | 2.26 | 2.36e-02 |
| $EE_{vgll3}EE_{akap11}$ | 3.66 | 0.58 | 6.27 | 3.59e-10 |
| Mean-centred length | -0.09 | 0.01 | -9.06 | 1.26e-19 |

Supplementary Table 7. Coefficients of the model comparing the effects of $EL_{vgll3}EL_{akap11}$ and $LL_{vgll3}EL_{akap11}$ genotypes on the odds of maturation. Genotype estimate is relative to the $LL_{vgll3}EL_{akap11}$ genotype.

| Coefficient | Estimate | SE | z-value | p-value |
|---|---|---|---|---|
| Intercept | -8.37 | 1.05 | -8.00 | 1.23e-15 |
| $EL_{vgll3}EL_{akap11}$ | 2.52 | 0.56 | 4.51 | 6.36e-06 |
| Mean-centred length | -0.11 | 0.01 | -7.23 | 4.85e-13 |
