## Supplementary Figure for "Refining the genomic location of SNP variation affecting Atlantic salmon maturation timing at a key large-effect locus"

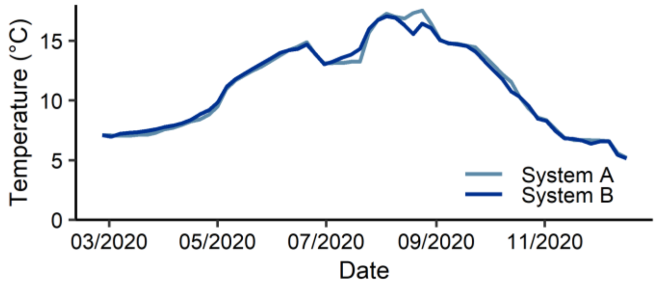


Supplementary Figure 1. Rolling 5-day average temperature for each water system for the post-hatch period of the experiment. Deviation between systems during late August was due to a temporary adjustment of water inflow in order to regulate oxygen levels.
